# Automated Detection of Records in Biological Sequence Databases that are Inconsistent with the Literature

**DOI:** 10.1101/101246

**Authors:** Mohamed Reda Bouadjenek, Karin Verspoor, Justin Zobel

**Affiliations:** Department of Computing and Information Systems, The University of Melbourne, Parkville, 3053, Australia

**Keywords:** Data Analysis, Data Quality, Bioinformatics Databases, Anomaly Detection

## Abstract

We investigate and analyse the data quality of nucleotide sequence databases with the objective of automatic detection of data anomalies and suspicious records. Specifically, we demonstrate that the published literature associated with each data record can be used to automatically evaluate its quality, by cross-checking the consistency of the key content of the database record with the referenced publications. Focusing on GenBank, we describe a set of quality indicators based on the relevance paradigm of information retrieval (IR). Then, we use these quality indicators to train an anomaly detection algorithm to classify records as *“confident”* or *“suspicious”*.

Our experiments on the PubMed Central collection show assessing the coherence between the literature and database records, through our algorithms, is an effective mechanism for assisting curators to perform data cleansing. Although fewer than 0.25% of the records in our data set are known to be faulty, we would expect that there are many more in GenBank that have not yet been identified. By automated comparison with literature they can be identified with a precision of up to 10% and a recall of up to 30%, while strongly outperforming several baselines. While these results leave substantial room for improvement, they reflect both the very imbalanced nature of the data, and the limited explicitly labelled data that is available. Overall, the obtained results show promise for the development of a new kind of approach to detecting low-quality and suspicious sequence records based on literature analysis and consistency. From a practical point of view, this will greatly help curators in identifying inconsistent records in large-scale sequence databases by highlighting records that are likely to be inconsistent with the literature.

## 1 Introduction

Bioinformatics sequence databases such as GenBank or UniProt contain large numbers of nucleic acid sequences and protein sequences. In 2017, GenBank alone contained over 228 billion nucleotide bases in more than 199 million sequences — a number that is growing at an exponential rate, doubling every 18 months.^1^ In commercial organizations, the primary reason for creating and maintaining such databases is their importance in the process of drug discovery, while in research they are used to understand the biological basis of disease. Thus, a high level of data quality is crucial.

However, since these databases are fed by direct submissions from individual laboratories and by bulk submissions from large-scale sequencing centers, they suffer from a range of data quality issues [1] including errors, redundancies, ambiguities, incompleteness, and as we will show, discrepancies such as inconsistency with the literature. Most of these records are linked to research articles in which the sequence was reported, but the need to manually create the records on such a large scale means that errors creep in and, given the volume, human curation alone is not sufficient for detection of these errors.

In this work, we seek to investigate and analyse the data quality of sequence databases from the perspective of a curator, who must detect anomalous and suspicious records. In contrast to previous research, which has concerned detection of duplicate records [2, 3, 4] and erroneous annotations [5, 6, 7], we emphasize detection of low-quality records that we define as being inconsistent with the published literature. Specifically, we propose that the literature that is linked to records in their “reference” fields be automatically used as background knowledge to check their quality. We explore a combination of information retrieval (IR) and machine learning techniques to identify records that are anomalous and thus merit analysis by a curator.

To provide insight into the data quality of the nucleotide records cited by articles available in PubMed Central^2^ (PMC) from a literature consistency point of view, we analyzed these records as illustrated in Figure 1. This figure shows the term overlap similarity^3^ between the record definition and different sections of its associated article(s) (representing the title, abstract, body, and the full text). There are three notable trends here: first, term overlap increases from title to body and full text since the size grows accordingly; second, there is a high term overlap of roughly 80% between the record description field and the literature body section; and third, when considering the overlap similarity between the description field of the records and the full text of their associated articles, for a small number of records in which the overlap similarity is below the value of the minimum whisker (0.4), there is low overlap or no overlap at all, thus statistically suggesting a data quality problem.

**Figure 1:**
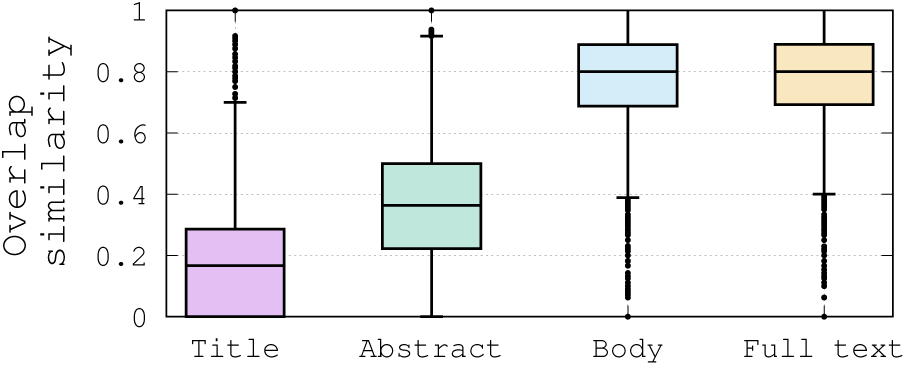
Overlap similarity between a record definition field and different sections of its associated document.

As an example, the record with accession number KM403369^4^ doesn’t share any terms with the article PMC4465667^5^ that is supposed to report on that record. Compared to the median value, which is roughly 80% similarity between a record description field and the body section of the article (see Figure 1), this association can be considered an outlier from a statistical perspective, and can be argued to be weak. While this observation is purely statistical, it may be an indicator of a low confidence in that record. Although this record is not necessary faulty, its characteristics in relation to the overall statistical distribution clearly suggest that it should be flagged as “suspicious”, and should be sent to a curator for further investigation.

Usually, a suspicious record is reported manually, by a curator whose the job consists mainly to check the database records, the record’s original submitter, or a third person who may use the database and notice the inconsistency of that record. To illustrate the difficulty of the task of identifying failing records, we analysed the distribution of record ages, for records which have been removed. This analysis showed that removed records have an average age of about 1 month at their removal time. This leads us to make two hypotheses: either (i) it takes about one month for a problematic record to be detected, or (ii) curators focus only on new records, while neglecting older ones. Either way, it is clear that there is a time window of only 1 month during which curators act. Hence, if a suspicious record is not identified in this time frame, it has a low probability of being spotted. These observations show the difficulty of the curator’s job, and the need for the development of automatic methods to assist them.

With the aim of assisting curators, and while focusing on GenBank, we present in this paper a method for detection of suspicious records based on their associated articles and also on the collection of articles as a whole. To the best of our knowledge, this work is the first to use the literature for data quality assessment of bioinformatics sequence databases. The contributions of this paper are as follows:

- We demonstrate that the research literature can be automatically used for assessing the quality of a record.
- We propose a list of quality indicators that correlate with the quality of a record. The quality indicators are then used to train a learning anomaly detection algorithm.
- Our experiments on the full PubMed Central collection show that, although less than 0.25% of the records in our data set are faulty, by automated comparison with literature they can be identified with a precision of up to 10% and a recall of up to 30%, while greatly outperforming the best baseline.

## 2 Related Work

There is a substantial body of research related to data quality in bioinformatics databases. Previous research has focused mainly on duplicate record detection and erroneous annotations, as reviewed below.

### 2.1. Duplicate records

Koh et al. [4] use association rule mining to check for duplicate records with per-field exact, edit distance, or BLAST sequence [8] alignment matching. Drawbacks of this method, and its poor performance, have been shown by Chen et al. [2, 3]. Similarly, Apiletti et al. [9] proposed extraction of association rules among attribute values to find causality relationships among them. By analysing the support and confidence of each rule, the method can show the presence of erroneous data. Other approaches also use approximate string matching to compute metadata similarity [10, 11, 12]. However, as they focus only on metadata, the underlying interpretation is that duplicates are assumed to have high metadata similarity, or that their sequences are identical.

Other approaches consider duplicates at the sequence level; they examine sequence similarity and use a similarity threshold to identify duplicates. For example, Holm and Sander [13] identified pairs of records with over 90% mutual sequence identity. Heuristics have been used in some of these methods to skip unnecessary pairwise comparisons, thus improving the efficiency. Li and Godzik [14] proposed CD-HIT, a fast sequence clustering method that uses heuristics to estimate the anticipated sequence identity and will skip the sequence alignment if the pair is expected to have low identity. Recently, Zorita et al. [15] proposed Star Code to detect duplicate sequences, which uses the edit distance as a threshold and will skip pairs exceeding the threshold. Such methods are valuable for this task, but do not address the problem of consistency or anomaly.

### 2.2. Erroneous annotations

Sequence databases exist as a resource for biomedicine, but the utility of the sequence of an organism depends on the quality of its annotations [12]. The annotations indicate the locations of genes and the coding regions in a sequence, and indicate what those genes do. That is, annotations serve as a reading guide to a sequence, which makes the scientific community highly reliant on this information. Although the research and development of algorithms for identifying coding sequences (CDSs) is still an active area in bioinformatics research, genome annotation has evolved greatly during past few years [16, 17, 18, 19]. However, the functional annotation of CDSs is particularly difficult to automate [20]. Current state-of-theart functional annotation methods integrate multiple types of evidences [21, 22, 23], but unfortunately the quality of functional annotations remains generally poor [24, 25, 26] and is highly dependent on resource-intensive manual curation [27, 28].

Previous research work on function annotation identified potential problems with large-scale annotation efforts [5, 29, 30], and mis-annotation is a growing concern among the general research community, as misannotations can have a several impacts in diverse biological areas [31, 32, 33]. Even in very small bacterial genomes, many misannotations may arise [34]. As for high-throughput functional annotations, errors may occur due to a variety of factors [35, 36], but the most common errors are over-annotations, in which a gene is given a specific but incorrect function [26, 34, 37]. Once made, functional annotation errors can be difficult to correct in large scale sequence databases and as functional annotations are often inferred from sequence similarity to other annotated sequences, errors may “propagate” to newly sequenced genomes through “(mis)annotation transfer” [38, 39, 7].

Overall, existing data quality analysis methods for sequence databases focus only on the internal characteristics of records. Our work demonstrates that the literature associated to records is a valuable external source of information for assessing the quality of sequence database records.

## 3 Background and Problem Definition

In this section, we first provide an overview of GenBank, the most commonly used sequence database, and we describe the structure of a sequence record in GenBank. Next, we discuss several kinds of data issues in bioinformatics sequence databases, and finally, we define in detail the problem we study.

### 3.1. GenBank overview

In this work, we mainly focus on GenBank as it is the most important and most influential sequence archive repository for research in almost all biological fields, whose data are accessed and cited by millions of researchers around the world. GenBank is produced and maintained by the National Center for Biotechnology Information (NCBI), and is considered to be an archive rather than a database, because multiple versions of a given record may be maintained for historical purposes. The sequence submission to GenBank can occur through: (i) direct submissions from scientists using BankIt^6^, which is a Web-based form, or the stand-alone submission program, Sequin^7^, or (ii) bulk submissions most often done by large-scale sequencing centers, which include Expressed Sequence Tag (EST), Sequence-tagged site (STS), Genome Survey Sequence (GSS), and High-Throughput Genome Sequence (HTGS). Upon receipt of a sequence submission, an accession number is assigned to the sequence, and then, it is released to the public database, where the entry is retrievable using Entrez^8^.

Due to the fact that records can be submitted by multiple research actors without any particular data quality control, errors may occur. Errors can seriously hamper the efficacy of analysis, data mining, and machine learning algorithms. Hence, a faulty record is usually reported manually, by a database curator, the record’s original submitter, or a third person who may use the database and notice the inconsistency of that record. However, updates and revisions of a GenBank sequence can also be made by the submitters at any time.

In addition to GenBank which can be considered as an unreviewed repository and thus may contain low quality sequences, NCBI also maintains other curated sequence databases such as the Reference Sequence (RefSeq). RefSeq provides a comprehensive, integrated, non-redundant, well-annotated set of sequences, including genomic DNA, transcripts, and proteins. RefSeq genomes are copies of selected assembled genomes available in GenBank, which have been generated by several processes including manual curation, which is known to be a tedious and painful task.

### 3.2. GenBank sequence record structure

The format of a sequence record can be regarded as having three parts: the header, which contains the information that applies to the whole record; the features, which are the annotations on the sequence; and the sequence itself. The header section is composed of several fields:

- *LOCUS* field: contains a number of different data elements, including locus name, sequence length, molecule type, and modification date. The locus name is designed to help group entries with similar sequences: the first three characters usually designate the organism; the fourth and fifth characters can be used to show other group designations, such as gene product; for segmented entries the last character is one of a series of sequential integers;
- *DEFINITION* field: a brief description of sequence or sequence’s function;
- *ACCESSION* field: a unique identifier for the record;
- *SOURCE* field: gives information about the sequence’s organism;
- *REFERENCE* field: lists a set of publications by the authors of the sequence that discuss the data reported in the record.

It is clear that the header part represents in itself a rich source of information.

Based on the fact that articles discuss the data reported in the records, and that there is high term overlap between the record definition and its associated articles as reported in Figure 1, we will primarily focus on the record definitions to assess the quality of the records from a literature consistency point of view.

### 3.3. Classification of Biological Data Quality Issues

Given a sequence record with its multiple data elements, the complex sequence submission process and the data integration pipeline defined to exchange data with other sequence databases, data quality issues may have physical or conceptual sources. Hence, Koh et al. [40, 1] proposed to classify biological data issues according to their presence in data items at mainly four levels of detail — individual attributes, individual records, individual databases, and multiple databases, as shown in Figure 2. Below, we briefly discuss these data quality issues.

**Figure 2:**
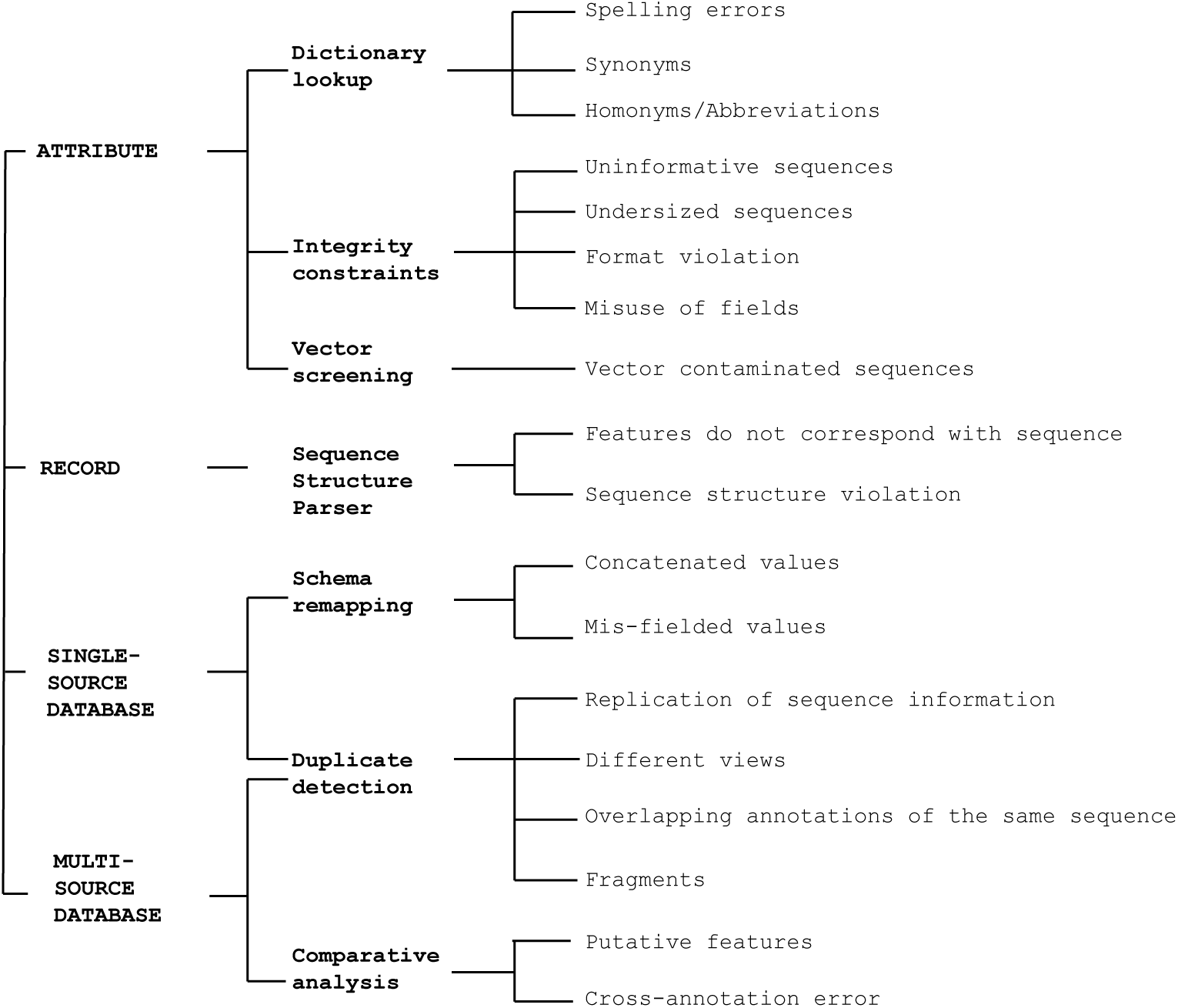
Classification of biological data issues by Koh et al. [40, 1].

#### 3.3.1 Attribute-level data quality issues

Attribute level data issues are field values with uninformative, invalid, erroneous or ambiguous content. Koh et al. [1] observed four main types of attribute level data quality issues — invalid attribute values, ambiguous attribute values, dubious sequences, and contaminated sequences.

##### Invalid values

Header data issues result from the use of non-standard names, free-text entries, and from the diversity of database schema used in different databases. The header information is usually entered and provided by the person who submits the original record to the database (such as the description). Hence, the header information may contain spelling errors or invalid field values. Koh et al. [1] identified 569 possible misspelled words affecting up to 20,505 nucleotide records in Entrez. For example, “immunoglobulin” is misspelled as “immunoglobin” in the record with accession number AB122023^9^. Another example is the organism “brachydanio rerio” (zebrafish) which is misspelled “brachidanio rerio” as in the record with accession numbers L25057^10^.

##### Ambiguity

As there is a lack of standardized naming conventions and controlled vocabulary use, vastly different definitions may be used in database records to refer to the same sequence. The naming errors include use of different names for the same sequence (synonym problem) or the same name for different sequences (homonym problem) [41]. For example, the scorpion neurotoxin “BmK-X” precursor has many possibly synonymous permutations. It is also known as “BmKX”, “BmK10”, “BmK-M10”, “Bmk M10”, “Neurotoxin M10”, “Alpha-neurotoxin TX9”, and “BmKalphaTx9”^11^.

Another type of error is the use of abbreviations, which may result in ambiguities. For example, the abbreviation BMK stands for “Big Map Kinase”, “B-cell/myeloid kinase”, “bovine midkine”, as well as for “Bradykinin-potentiating peptide”. GK is the abbreviation for both “Glycerol Kinase” and “Geko” gene of Drosophila melanogaster (Fruit fly).

In free-text fields, a wrong piece of information can be entered as field value. For example, the description of the sequence with the accession number AP011615^12^ is “Arthrospira platensis NIES-39, *** SEQUENCING IN PROGRESS ***, 19 ordered pieces”.

##### Dubious sequences

The sequence is represented as a string of letters denoting the 20 amino acids in the case of a protein sequence and the 4 nucleotides in the case of nucleotide sequence. Each base or residue is thus limited to it alphabetical representation and “X” is used to denote an unknown residue, and “N” to denote an unknown base. A base or residue which doesn’t correspond to its set of special letters is invalid and can be caused by an erroneous data entry. For example, the sequence with the accession number AC000016^13^ contains 11% of unknown bases.

A sequence may also contain invalid symbols for nucleotides or amino acids, or it can be shorter than its logical size. The length of protein sequences usually ranges from 6 to a few thousands residues. However, Koh et al. [1] found 3,327 undersized protein sequences which are shorter than six residues in the public databases using Entrez (as of Sep 2004), among which 1,887 contain only one residue.

##### Contaminated sequences

There are cases where a DNA sequence contains vectors used for cloning; vector-contaminated sequences may be submitted to the database. Vectors are agents that carry DNA fragments into a host cell. The vector sequences probe and bind the DNA fragments at the 5’ and 3’ sites. The DNA fragment is then isolated from its vectors by cutting at the restriction enzyme sites. The existence of vector-contaminated sequences was first reported in 1992; 0.23% of 20,000 eukaryotic entries were found to be contaminated [42]. In 1999, [43] reported that up to 0.36% or 3,029 of the sequences in GenBank contain contamination of the cloning vectors.

#### 3.3.2. Record-level data quality issues

Conflicting information exists in the single record among two or more attributes — Koh et al. [40] call them the record-level data quality issues. Two types of record-level data quality issue are found in sequence records: sequence structure violations and inconsistent content with related references.

##### Sequence Structure Violations

It is known that a gene structure has a set of logical constraints, and any infringement of these constraints constitutes a possible feature issue. Such logical constraints include that introns and exons must be non-overlapping except in cases of alternative splicing. Koh et al. [1] observed that 12 out of 42,359 nucleotide sequences had overlapping intron/exon regions. For example, Syn7 gene of putative polyketide synthase in NCBI TPA record BN000507^14^ has overlapping intron 5 and exon 6. The rpb7+ RNA polymerase II subunit in GenBank record AF055916^15^ has overlapping exon 1 and exon 2.

##### Inconsistent with the literature

Usually, each record is associated with a list of publications by the authors of the sequence that discuss the data reported in the record. However, it is possible that a record is inconsistent with the information provided in the literature in general, and in the articles related to that record in particular.

For example, in the study of Dengue virus, Koh et al. [40] observed mis-annotations in Swiss-Prot record P27915^16^ and PIR record GNWVD3 [44]. The NS1/NS2A and NS4A/NS4B junctions given in these Dengue type 3 complete RNA sequences did not match the regions given in the reference of these records [45]. While manual checking of such inconsistencies by cross-referencing the database content with their corresponding literature is tedious, computational detection of discrepancies of the sequence annotations with its references is also non-trivial and may require complex text-mining solutions.

#### 3.3.3. Single Database level data quality issues

##### Annotation errors

The features of a sequence are often directly submitted by the author of the sequence. The features can be derived experimentally or inferred. Computationally inferred features are usually based on sequence homologues and are derived using annotation tools. Hence, multiple database records of the same nucleotide or protein sequences may contain inconsistent or conflicting feature annotations. Koh et al. [1] refer to such data issues as cross-annotation errors. They identify possible causes of cross-annotation errors as: (i) different expert interpretations, (ii) mis-annotation of sequence function, and (iii) inference of features or annotation transfer based on best matches of low sequence similarity.

Annotation errors commonly result from mis-annotation or from data entry errors. In GenBank entries that contain splice site features in Arabidopsis thaliana, some 15% were found to have incorrect annotations [46]. Another study [39] found that 24% of the Chlamydia trachomatis sequences contained erroneous functional assignments. Another form of annotation error is caused by inaccurate inference of features from homologues.

##### Sequence duplication

Sequence duplication is also observed in sequence databases [2]. There are three types of redundancy: (1) the same sequence and annotations can be found in multiple records, (2) the same sequence but different annotations are found in multiple records, and (3) partially overlapping annotations of the same sequence exist in multiple databases which have different data views. For example, the records AAG39642^17^ and AAG39643^18^ contain identical sequences with exactly the same annotations.

#### 3.3.4. Multiple Database level data quality issues

Due to the existence of heterogeneous database schema, massive data transformation is carried out in the databases during large-scale uploads or during data exchange. The transformation of data records from one schema to another may cause data integration problems, where data may be mapped to the wrong fields.

#### 3.3.5. Summary

In this work, we are interested in the detection of records that contain errors and inconsistencies through the analysis of the header section, and through a cross validation with the published literature. Thus, the model we built will not be able to detect errors related to the features or the sequence itself such as contaminations, undersized sequences, cross annotation errors, etc. Rather, we focus exclusively on detecting inconsistencies with the published literature as discussed in Section 3.3.2. While this represents only a portion of the potential errors, the method will be shown to add value to the broader task.

### 3.4. Research problem statement

We propose to follow an IR approach, where a database record is regarded as a query, and its associated articles as the relevant documents. We use the term “query” to refer to a record, and the term “relevant documents” to refer to the set of its associated articles in its reference field.

We define the problem we study in this paper as follows. Given:

- a collection of documents which represents the domain literature knowledge *D* =< *d*_1_*, d*_2_*,…, d*_*n*_ *>*
- a set of annotated queries *Q* =< (*q*_1_*, y*_1_), (*q*_2_*, y*_2_)*,…,* (*q*_*m*_*, y*_*m*_) *>*, where *y*_*m*_ ∈ {confident, suspicious}
- the set of relevant documents that meet the information need of each query *D*_*R*_ =*<D*_*R*_1__,*D*_*R*_2__,…, *D*_*R*_*m*__*>*

we aim to retrieve and identify queries that are not consistent with their relevant documents or with the collection as whole, indicating that their description in the database record is incompatible with the information given in the corresponding publication, and which can therefore be flagged as “*suspicious*”. The resulting tool is expected to be used at curation time, and should send such “*suspicious*” records to curators for review.

## 4 Quality Factors

In this section, we introduce the features that we will consider as record quality indicators. We describe two kinds of features: record-based features, which mainly focus on the characteristics of the records; and IR-based features, which mainly focus on query quality predictors. Our approach here is to define a wide variety of features and then identify which of them are most valuable in classification.

### 4.1 Record-based features (9 features)

The characteristics of a record are strong indicators of its quality. We define several features that consider a record as a whole. Hence, we mainly rely on basic and intuitive quality factors, such as the record popularity, as well as building on recommendations given by the International Nucleotide Sequence Database Collaboration (INSDC) for the record structure.

#### Organism popularity (1 feature)

Based on the intuition that organisms that have rarely been sequenced and deposited in a sequence database are more likely to have suspicious records, we consider the popularity of the main organism of a record as a quality feature. We define the popularity of an organism as the number of records that relate to that organism divided by the total number of records.

#### Record definition structure (3 features)

The INSDC suggests that the record definition should have the following specific format:^19^ (i) it should start with the common name of the source organism; (ii) it gives the criteria by which this sequence is distinguished from the remainder of the source genome, such as the gene name and what it codes for, or the protein name and mRNA, or some description of the sequence’s function (if the sequence is non-coding); (iii) if the sequence has a coding region, the description may be followed by a completeness qualifier, such as ‘cds’ (complete coding sequence). We define boolean attributes to indicate whether each of these rules is respected in a record or not.

#### Record popularity (1 feature)

a popular sequence record is more expected to have been checked by other users, and hence be a confident record. Thus, we include the popularity as a quality feature and define it simply as the number of citations the record has.

#### Coding sequence (3 features)

For a sequence with a coding region, the coding sequence (CDS) field in the features section of the records is one of the most important fields. Based on the feature table format designed jointly by GenBank, the EMBL Nucleotide Sequence Data Library, and the DNA Data Bank of Japan,^20^ the CDS field should specify: (i) the region of nucleotides that corresponds with the sequence of amino acids in a protein (location including start and stop codons), (ii) the gene name, and (iii) the product/protein name. For each CDS field of a record, we check its validity (i) by ensuring that the CDS region is within the sequence range, (ii) by ensuring that the gene name is valid and is given into the annotations of Gene Ontology (GO) ([47]), and (iii) by ensuring that the gene exists and is described in the NCBI gene database [48]^21^. Hence, we define three quality attributes for (i), (ii), and (iii) based on an aggregation of all CDS fields of each record.

#### Definition length (1 feature)

the length of the definition may indicate the degree of precision given to describe a record. Hence, we include the length as a quality factor, and we define it as the total number of terms.

### 4.2 IR-based features (203 features)

To find indicators or features to represent the quality of each query (record), we draw on the large body of previous work on query quality prediction ([49, 50, 51]). While some of these features such as Overlap Similarity are stand-alone, other features such as Average Term Frequency are derived from term level statistics ([51]). These include predicting the quality of queries using either pre-retrieval indicators like Query Scope, that is, they are calculated for a query as a whole, or post-retrieval indicators like Query Clarity, that is, they involve performing an initial retrieval and hence are more expensive to compute. We describe the set of query quality predictors we used. As stated previously, to compute these IR-based features, we consider the record *definition* field as the query.

#### Query clarity (QC) (18 features)

Developed by [49], this post-retrieval factor is the Kullback-Leibler divergence of the query model from the collection model. QC is computed as:

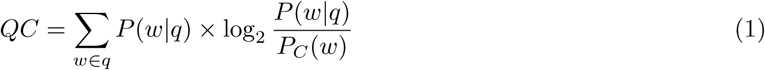

where *P* (*w|q*) is the probability of the occurrence of the word *w* in the query model, and *P*_*C*_ (*w*) is the probability of the occurrence of *w* in the collection. The query model is estimated from the top-*k* ranked documents retrieved after an initial run of the original query. We computed different QC scores based on different configuration options *k* ∈ {1, 5, 10, 20, 50, 100} × *w* ∈ {title, abstract, body}.

#### Simplified clarity score (SCS) (3 features)

To avoid the expensive computation of query clarity, [52] proposed simplified clarity score as a comparable pre-retrieval performance factor. It is calculated as:

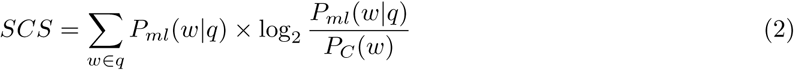

where *P*_*ml*_(*w|q*) is the probability of the occurrence of the word *w* in the query. We also computed SCS based on different configuration options of *word ∈ {*title, abstract, body}.

#### Relevant-documents clarity score (RDCS) (3 features)

We also propose to compute the clarity score based on a query model estimated from the relevant documents themselves, while considering separately three different fields of the relevant documents {title, abstract, body}.

#### IDF-based features (24 features)

We calculate the *Inverse Document Frequency* (IDF) of each query term *w* as:

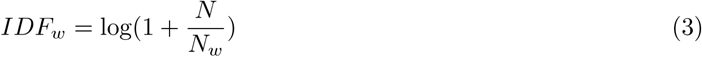

where *N*_*w*_ is the document frequency of *w* while considering separately three different fields {title, abstract, body}, and *N* is the number of documents in the collection. For each query we calculate the sum, standard deviation, minimum, maximum, arithmetic mean, geometric mean, harmonic mean, and coefficient of variation of the IDFs of constituent terms.

#### TF-based features (24 features)

We calculate the *Term Frequency* (TF) of each query term *w* in a relevant document *d* as:

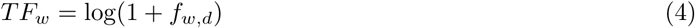

where *f*_*w,d*_ is the number of time *w* occurs in *d* while considering separately three different fields {title, abstract, body}. For each *TF* value of each term, we calculate aggregate values similar to those for IDF as quality factors.

#### Similarity of collection–query (SCQ) (24 features)

Proposed by [53], this query quality factor is based on the hypothesis that queries that have higher similarity to the collection as a whole will be of higher quality. For each term *w* in the query, SCQ is computed as:

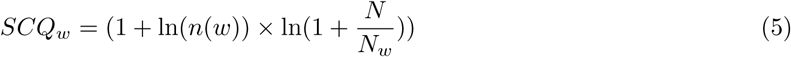

where *n*(*w*) is the frequency of the term *w* in the collection while also considering separately three different fields {title, abstract, body}. Based on the SCQ values of each term, we also calculated aggregate values similar to those for IDF.

#### Inverse collection term frequency features (ICTF) (24 features)

The inverse collection term frequency of a term *w* is defined as:

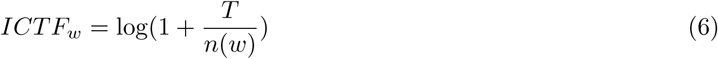

where *n*(*w*) is the frequency of the term *w* in the collection and *T* is the number of term occurrences in the collection while considering separately three different fields {title, abstract, body}. Using the ICTF values, we calculate aggregate statistics similar to those for IDF.

#### Query scope (QS) (4 features)

Query scope ([52]) is a measure of the size of the retrieved document set relative to the size of the collection. We can expect that high values of query scope are predictive of poor-quality queries as they retrieve far too many documents. QS is computed as:

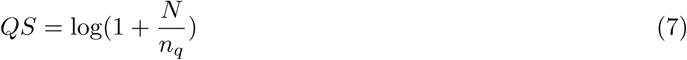

where *n*_*q*_ is the number of documents that match the query terms while considering separately four different fields {title, abstract, body, all document}.

#### Similarity of relevant documents–query features (48 features)

Based on the fact that a high similarity value between a query and its relevant documents reflects a high query quality, we include several information-theoretic, statistical, and practical similarity measures as quality indicators. These similarity measures are: matching, overlap, Jaccard, Dice, cosine, mutual information (MI), and Okapi BM25. We also used various IR similarity ranking functions including: the sum of TFIDF scores (SumTFIDF), the Lucene vector-space model score (LuceneVSM),^22^ the BM25 score ([54]), language model scores based on (i) the Jelinek-Mercer smoothing (LMJelinekMercer) ([55]) and on (ii) a Bayesian smoothing using Dirichlet priors (LMDirichlet) ([55]), and an information-based score (IBSimilarity) ([56]). These similarities are computed while considering separately four different fields {title, abstract, body, all document}.

#### Retrieval performance score (RPS) (28 features)

Based on the relevance paradigm of IR, we assume that a good quality record should rank its corresponding articles highly. Thus, we use the reciprocal rank evaluation measure to define the RPS as follows:

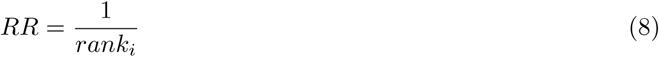

where *rank_i_* is the rank of the first relevant document in the retrieved list of documents that matches the query *q*_*i*_ returned by the system. We have also considered query expansion using the following terms related to the organism: (i) scientific name, (ii) common names, (iii) synonyms, (iv) abbreviations, (v) misnomers, and (vi) misspellings. These terms are extracted from the NCBI Taxonomy,^23^ which is a curated classification and nomenclature for all of the organisms in the public sequence databases. The basic intuition is that: if (i), (ii), (iii), and (iv) improve the retrieval performance, there is a mismatch between the record and its corresponding article, and thus, the record may be of low quality. Also, if (v) and (vi) improve the retrieval performance, the article is clearly reporting the record using incorrect terms, and thus, the record is probably of low quality. Here we consider querying separately four different fields {title, abstract, body, all document}.

#### Citation of the main organism in the article (3 features)

The citation of the main organism in the relevant document is an important quality factor since the article is supposed to report the content of the record. Hence, we include this quality factor a binary feature while considering separately four different fields {title, abstract, body}.

In total, we have defined 9 record-based features and 203 IR-based features, for a total of 212 features that characterize the quality records.

## 5 Experimental Setup

In this section, we describe the dataset we have constructed from publicly available resources, and then introduce the learning algorithm we used to classify records as *“confident”* or *“suspicious”*.

### 5.1 Data description

Now we provide details of the dataset that we evaluate in this paper.

#### Articles

We used the PubMed Central Open Access collection^24^ (OA), which is a free full-text archive of biomedical and life sciences journal literature at the U.S. National Institutes of Health’s National Library of Medicine. The release of PMC OA we used contains roughly 1.13 million articles, which are provided in an XML format with specific fields corresponding to each section or subsection in the article. We used the Lucene IR System^25^ to index the collection, with the default settings for stemming and English stop-word removal. We defined a list of biomedical keywords, which should not be stemmed or considered as stop-words, such as the protein names *“THE”* and *“Is”*. Each section of an article (title, abstract, body) is indexed separately, so that different sections can be used and queried separately to compute the quality features.

#### Sequences

We work with the GenBank nucleotide database, but limit the sequence database records we work with to those that are cited by the PMC OA article collection. Specifically, we used a regular expression to extract GenBank accession numbers mentioned in the PMC OA articles, thereby identifying literature that refers to at least one GenBank identifier. This resulted in a list of 733,779 putative accession numbers. Of these, 494,142 were valid GenBank nucleotide records that we were able to download using the e-utilities API ([57]).^26^ Among the valid records, only 162,913 records also cite the corresponding articles (as determined by matching their titles). This process gave us a list of 162,913 pairs of record accession numbers and PMC article identifiers, which cite each other. Note that for the 331,229 records that we have put aside, each record cites an article; however, we do not have access to all articles through PMC OA.

Each record in this dataset was labelled as *“alive”* or *“dead”*, an attribute that we obtained using the eSummary API ([57]). Note that the records that are reported as *“dead”* are explicitly labelled as such in GenBank. For example, the record with the accession number DW407270^27^ is explicitly reported as *“dead”* as it is currently removed from GenBank. However, records that are *“alive”* are implicitly labelled by not being dead. In the classification task, we consider dead records to be *“suspicious”* and all other records to be *“confident”* in our labelling. We acknowledge that an *“alive”* record does not necessarily indicate that it is of good quality. However, we made this assumption motivated by the fact that, overall, the records are of good quality, whereas only a small fraction of the data may be faulty. The set of unidentified faulty records can be regarded as *“zombie”* records. This has been observed and reported in [58], where the authors carried out a biocuration task. The authors randomly selected a sample of 100 alive records in the dataset, and a biocurator manually found 5 records to be faulty (*“zombie”* records). Although the sample of records was small^28^, we believe that the rate of faulty records in the whole dataset is of the order of roughly 5%. We rather give this number to support our decision to let this small fraction of *“zombie”* records to be in our dataset, as they would have an insignificant impact on the learned model.

We provide more detailed statistics on our final dataset in Table 1. For example, an article cites 0.51 records on average; the article PMC2848993^29^ cites 8,062 records. A record is cited on average 1.17 times, while the record A23187^30^ is the most cited. Among the 162,913 records for which the relevant articles are in the PMC OA dataset, 162,486 are alive and only 427 are dead. Hence, our dataset is skewed toward negative examples (alive records) with only a few positive examples (dead records).

**Table 1:** Dataset Statistics.

| Articles statistics |  |  |  |  |
| --- | --- | --- | --- | --- |
|  | Citations of records |  |  |  |
| # articles | Avg | Median | Max | Max entity |
| 1,135,611 | 0.5114 | 1 | 8,062 | PMC2848993 |
| Sequence statistics |  |  |  |  |
|  | Record citations |  |  |  |
| # articles | Avg | Median | Max | Max entity |
| 494,142 | 1.1755 | 1 | 783 | A23187 |
| Records for which the relevant article is in our PMC OA dataset |  |  |  |  |
| #records | Alive records |  | Dead records |  |
| 162,913 | 162,486 |  | 427 |  |
| Organism taxonomy statistics |  |  |  |  |
| 77,632 genus and 1,118,198 species |  |  |  |  |
| Class name |  | Total | Avg | Max |
| Scientific names |  | 1,195,830 | 1 | 1 |
| Synonyms |  | 165,867 | 0.1387 | 55 |
| Misspellings |  | 24,776 | 0.0207 | 14 |
| Common names |  | 12,820 | 0.0107 | 10 |

#### Organism taxonomy

To gather more information about the record organisms, such as the list of synonyms, acronyms and common names, we used the NCBI Taxonomy database. This is a curated classification of all of the organisms in the public sequence databases. The NCBI taxonomy contains and describes of roughly 10% of the existing species of life on the planet.

### 5.2 Anomaly detection algorithm

Given as input a set of quality indicators for each record, our goal is to combine these inputs to produce a value indicating whether the record is *“confident”* or *“suspicious”*. To accomplish this, we used the Support Vector Machines (SVM) classification algorithm ([59]), which is one of the most widely-used and effective classification algorithm.

Each record *m* is represented by its vector of *k* quality indicators *x*_*m*_=[*x*_*m*1_*, x*_*m*2_*,…, x*_*mk*_] and its associated label *y*_*m*_ ∈ {confident, suspicious}. We used the SVM implementation available in the LibSVM ([60]) package. Both Linear and RBF kernels were considered in our experiments. The regularization parameter *C* (the trade-off between training error and margin), the gamma parameter of the RBF kernel, and the penalty parameter *w*_*i*_ that penalizes negative examples due to the skewed nature of the dataset were selected from a search within the discrete sets {10^−5^, 10^−3^*,…,* 10^13^, 10^15^}, {10^−15^, 10^−13^*,…,* 10^1^, 10^3^}, and {10, 20*,…,* 50, 100, 200} respectively, using 10 fold cross validation.

Although the differences were not substantial, experiments with the best RBF kernel parameters performed slightly better than the best linear kernel parameters for the majority of the validation experiments. Unless otherwise noted, all presented results were obtained using an RBF kernel, with C set to 10^−3^, gamma set to 10^−3^, and *w*_*i*_ set to 100.

## 6 Experimental Evaluation

We now report and discuss the main results of the experimental evaluation, considering both the effectiveness of the method and our interpretation of which features are valuable in classification.

### 6.1 Feature analysis

To explore the relationship between features and the record quality labels, we undertook a feature analysis task. A general method for measuring the amount of information that a feature *x*_*k*_ provides w.r.t. predicting a class label *y* (*“confident”* or *“suspicious”*) is to calculate its mutual information (MI) *I*(*x*_*k*_*, y*) or Pearson’s chi-squared test *χ*^2^(*x*_*k*_*, y*). In Table 2, we present the list of ten top-ranked features using these two metrics. The two lists are roughly similar except for the tenth line, where MI introduces a record-based feature. These two lists led us to make the following observations:

**Table 2:** Ranking of the most important features using two different metrics.

| Rank | Mutal Information |  | Pearson’s chi-squared test |  |
| --- | --- | --- | --- | --- |
|  | Field | Feature | Field | Feature |
| 1 | title | LMDirichlet Score | title | LMDirichlet Score |
| 2 | abstract | SumTFIDF score | abstract | SumTFIDF Score |
| 3 | abstract | LMDirichlet Score | abstract | RDCS |
| 4 | abstract | BM25 Score | abstract | LMDirichlet Score |
| 5 | abstract | RDCS | abstract | BM25 Score |
| 6 | abstract | IBSimilarity Score | abstract | IBSimilarity Score |
| 7 | title | SumTFIDF Score | title | BM25 Score |
| 8 | title | BM25 Score | title | IBSimilarity Score |
| 9 | title | IBSimilarity Score | title | SumTFIDF Score |
| 10 | x | Popularity Organism | abstract | LMJelinekMercer Score |

1. Features based on the similarity between the relevant documents and the record are the most informative (8/10 for mutual information and 9/10 for chi-squared). This confirms that a good and a confident record is highly discussed in its associated articles.
2. IR similarity ranking functions are the most informative features. They take into account the information carried in both the query and the documents, in contrast to statistical similarity measures.
3. For both rankings, the top feature is a language-model similarity score. It computes the similarity between the record definition and the title of the relevant document, using Bayesian smoothing with Dirichlet priors. This shows that a confident record is one in which the description has a high probability of having been generated from the title of its associated document.
4. Top features are mainly based on short and medium document fields (that is, title and abstract). This reflects the fact that confident records can be expected to be referenced and discussed earlier in the article.
5. Almost all top features are IR-based features. Only one record-based feature appears in the two rankings (popularity of the organism in the MI ranking). This confirms our initial assumption that the literature is a strong resource for checking the quality of a record.
6. Finally, some features that we experimented with were identified as being entirely uninformative; the mutual information values obtained were null (0). This includes: the popularity of the organism, some scores based on the standard deviation, and some scores based on the coefficient of variation.

### 6.2 Classification performance

The effectiveness of our method (denoted SVM LBF+RBF, for SVM with Literature-based Features and Record-based Features), is summarized in Table 3, broken down by the two classes in the data. The last column of the table shows the harmonic mean between the F1-Scores of the two classes. We provide a comparison with two variants of this method and four baseline methods:

- SVM RBF: SVM classifier trained with only record-based features.
- SVM LBF: SVM classifier trained with only IR-based features.
- Majority class: a naïve approach that simply predicts the most common class in the data (that is, classify everything as *“confident”*).

**Table 3:**
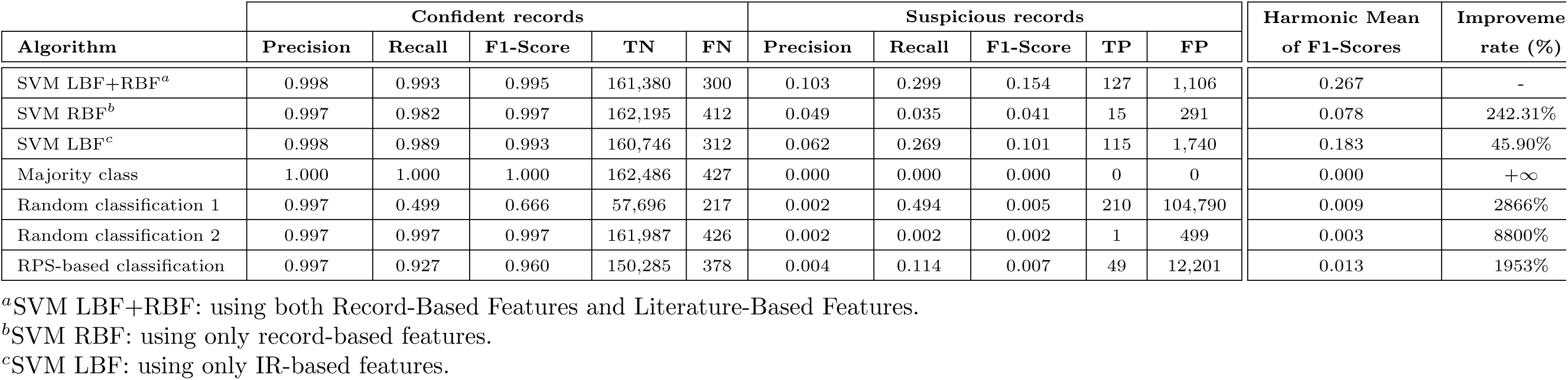
The results of the classification accuracy.

- Random classification 1: which classifies a record as *“confident”* or *“suspicious”* with a 50% probability. The classification was performed 1000 times independently over the full dataset; we report average results.
- Random classification 2: classifies a record as *“confident”* or *“suspicious”* with a 99.73% probability of being classified as confident. This value reflects the natural distribution of the source data, since 99.73% of the records are ‘alive’. The classification was also performed 1000 times independently over the full dataset; we report average results.
- RPS-based classification: classifies a record as *“confident”* or *“suspicious”* based on a fixed threshold (0.05) for the RPS value.

Our method (SVM LBF+RBF) shows a statistically significant improvement over the best baseline (relatively, 45.90% over SVM LBF). Second, the results confirm again our initial assumption that the literature is a strong support to assess the quality of records in sequence databases. Third, by comparing SVM RBF and SVM LBF, we conclude that the associated literature provides better evaluation of quality than can be obtained by examining only the records. This has also been shown in the previous section through the feature analysis. Due to the skewed nature of the dataset, all algorithms tend to classify the records as *“confident”*, which results in high precision and recall values for this class, but these results are not very informative. It is more meaningful to consider the performance over the class of *“suspicious”* records.

The instance-level TP and FP values given in Table 2 for each method illustrate how many FP records would need to be reviewed in order to find the small number of TPs in each case. This allows to compare the curator task difficulty for each method; the curator would need to examine everything retrieved (all Positives) and then make a decision as to whether it was a correct retrieval of a suspicious record (TP) or a perfectly valid record (FP from the perspective of *“suspicious”*). The table shows how much work needs to be done in each scenario, and demonstrates that our method considerably reduces the curation workload. In particular, to get reasonable recall with the random approach, the curator needs to review more than 100,000 records. Indeed, the curator would need to review 85x as many records with a random approach as compared to our method, for a gain of only 83 TP records (respectively: 1,233 vs. 105,000 positive records; 127 vs. 210 true positives identified). This highlights the substantial amount of effort saved using the approach we are proposing.

Table 3 shows relatively low values for precision and recall compared with some other machine learning problems. However, first, the dataset is highly imbalanced, with far more records labelled as *“confident”* than labelled as *“suspicious”*. Second, the records that are labelled as *“suspicious”* have been explicitly labelled, whereas records that are *“alive”* are implicitly labelled and assumed (perhaps wrongly) to be *“confident”*; an alive record may be a low-quality record that was missed in error. Hence, training a learning algorithm on unlabelled data leads to poor effectiveness, particularly when the minority class is the most likely class to be missed. Probably some of the records which have been classified as *“suspicious”* by our learning algorithm are *“zombie”* records; meaning that they are labelled in the data set as *“confident”*, but are in fact problematic. We discuss below typical examples of records which have been incorrectly classified by our method (false positives and false negatives). The profile of each example is given in Table 4, using the top 7 features obtained in the feature analysis in Section 6.1.

**Table 4:** Example profiles.

|  |  | Example 1 | Example 2 | Example 3 | Example 4 |
| --- | --- | --- | --- | --- | --- |
|  | **Accession** | FJ824848 | CP006742 | BK008760 | DW407270 |
|  | **PMC** | PMC2675058 | PMC3923885 | PMC4233938 | PMC1621083 |
|  | **Type** | FN | FN | FN | FP |
| 1 | **Tit. LMDir.** | 0.00 | 0.00 | 1.4194 | 18.78 |
| 2 | **Abs. SumTFIDF** | 0.00 | 0.00 | 2.25 | 27.04 |
| 3 | **Abs. LMDir.** | 0.00 | 0.00 | 3.3450 | 30.26 |
| 4 | **Abs. BM25** | 0.00 | 0.00 | 2.80 | 65.00 |
| 5 | **Abs. RDCS** | 0.00 | 0.0302 | 0.034 | 0.68 |
| 6 | **Pop. Organism** | 0.00000613 | 0.00 | 2.4828 | 0.00086 |
| 7 | **RPS** | 0.00 | 0.0019 | 0.20 | 1.00 |

#### Example 1

(False Positive) The record with accession number FJ824848^31^ has been classified as *“suspicious”* by the algorithm, and presents the typical profile of a suspicious record as given in Table 4. This record presents the complete sequence of the cloning vector pDMK3.^32^ First, the record definition does not give much information. Second, this record presents an organism for which there are relatively few other records. Third, the content of the record is mentioned neither in the title of the article nor in the abstract of the article with PMC identifier PMC2675058^33^. By examining the content of the article, we have noted that this cloning vector is mentioned as pDMK2. This case leads us to make two inferences: either the title of the record is incorrect, or the article uses an incorrect term to refer to this cloning vector. Consequently, we contacted the corresponding author of PMC2675058, Anders Sjöstedt, who acknowledged the error by saying: “For practical purposes this doesn’t matter since the two vectors are identical with the exception of two additional restriction sites in pDMK3. We should have stated pDMK3 in the paper so it can be denoted as a typographical error.” (personal communication, Anders Sjöstedt). However, since both pDMK2 and pDMK3 exist, we believe that this error cannot be considered as typographical error, but rather as a confusion and inconsistency between that record and its associated article.

#### Example 2

(False Positive) The record with accession number CP006742^34^ has also been classified as *“suspicious”* by the algorithm, and also presents the typical profile of a suspicious record as given in Table 4. This record presents the complete genome of the Bacillus anthracis organism. In fact, according to the article that reports the content of that record (accession number PMC3923885^35^), this record is supposed to present the chromosome Cow1. However, this chromosome is not mentioned in the record. We have contacted the corresponding author, and he answered as follows: “After some reasoning, we decided to call the isolates cow 1,2,3.. etc in the paper to increase readability. Their names in our strain collection is however different. As far as I can see CP006742 is the B. anthracis chromosome sequence that we submitted, although I haven’t checked every basepair…” (personal communication, Bo Segerman). Hence, we believe that this little inconsistency between the record and its associated article has prompted the algorithm to classify this record as *“suspicious”*.

#### Example 3

(False Positive) The record with accession number BK008760^36^ has also been classified as *“suspicious”* by the algorithm. This record presents relatively low values in its profile compared to the false negative example presented as given in Table 4. According to the article that reports the content of that record (PMC identifier PMC4233938^37^), this article is supposed to show two genes, Atg8a and Atg8b. However, the record is only showing the coding sequence of the gene Atg8a. We also have contacted the paper’s authors who acknowledged the error by saying: “Thanks for your important notification. Indeed this is an entry error. The error was corrected in a Corrigendum that was published soon after the original paper” [61] (personal communication, Assaf Vardi). Although the error is again in the research article itself, this shows another record-literature inconsistency example, which illustrates a false positive that contributed to the low precision-recall values obtained.

#### Example 4

(False Negative) In contrast, the record with the accession number DW407270^38^ has been classified *“confident”* by our method, while its current status is *“dead”*. The record references a popular organism and is correctly discussed in its associated article. We did not discover any mismatch or conflict between this record and its associated article (PMC identifier PMC1621083^39^). In fact, this record has been removed because the underlying biological material suffered from bacterial contamination with pseudomonas fluorescens, an issue that could only be resolved upon consideration of the similarity of the record sequence to other sequences via a BLAST search. This example leads us to identify an important distinction — a record can be coherent from one perspective (literature consistency) while being inconsistent from another (biological content). It highlights the need to consider data quality from more than one perspective, and demonstrates that literature alone is insufficient to detect all suspicious records. It further explains the limitations on performance of our method.

The examples discussed above in this section are not necessarily faulty records. We have rather presented these examples to provide a brief justification for why we obtained low precision-recall values. This motivates us to build a manually curated dataset in order to remove any ambiguous and noisy records that may lead to the build of a biased model, and to develop features that will capture further aspects of sequence record quality. This will form part of our future work.

## 7 Conclusions and Future Work

In this paper we have introduced a list of factors that correlate with the quality of a record. We used these quality indicators to train an anomaly detection algorithm based on supervised learning to classify records as *“confident”* or *“suspicious”*. We then performed a complete analysis on the full PubMed Central collection. The main outcome of this work is evidence that, in addition to the sequence itself, the literature is a valuable external resource that can be used to assess the quality of a database record.

Despite the fact that our method significantly outperforms the suggested baselines, we obtained somewhat low effectiveness scores, compared to some other common machine learning problems. Therefore, we undertook a feature analysis and a failure analysis, examining specific cases that indicate causes of this performance — in particular, identifying that the ground truth may contain errors as well as recognising that literature alone is insufficient to represent the full spectrum of data quality issues.

This work is to the best of our knowledge the first use of the literature as a tool for addressing the data quality problem in biomedical sequence databases. We have shown that the approach can identify problematic records with enough accuracy to be of value to curators, potentially reducing the effort required to remove low-quality records by nearly two orders of magnitude.

Our current dataset relies on data from GenBank to obtain labels, and the negative labels are only derived implicitly. This suggests two directions for future work. First, it would be desirable to construct a manually curated dataset explicitly for development of automated quality analysis techniques. Second, there is a need for new unsupervised learning methods for anomaly detection. It may be, for example, that good and bad records have distinct distributions of attribute values, so that methods such as k-nearest neighbour or local outlier factor ([62]) could be applied. Having established that automated literature analysis can be applied in practice to this task, the challenge now is to improve performance and further reduce the effort needed to clean databases. We also expect that leveraging external textual information to support data cleaning will have broader application in other database contexts.

1 http://www.ncbi.nlm.nih.gov/GenBank/statistics/

2 http://www.ncbi.nlm.nih.gov/pmc/

3 We use the overlap similarity to emphasize the number of terms of a record definition that are in its associated article. Here, *Overlap* (*X*_1_ *, X*_2_).= *|X*_1_*∩ X*_2_ *|/min* (*|X*_1_*| |X*_2_ *|*)

4 http://www.ncbi.nlm.nih.gov/nuccore/KM403369

5 http://www.ncbi.nlm.nih.gov/pmc/articles/PMC4465667/

6 https://www.ncbi.nlm.nih.gov/WebSub/?tool=GenBank

7 https://www.ncbi.nlm.nih.gov/Sequin/

8 The Entrez Global Query Cross-Database Search System is a federated search engine, or web portal that allows users to search many discrete health sciences databases at the National Center for Biotechnology Information (NCBI) website.

9 http://www.ncbi.nlm.nih.gov/nuccore/AB122023

10 http://www.ncbi.nlm.nih.gov/nuccore/L25057

11 http://www.uniprot.org/uniprot/O61705

12 http://www.ncbi.nlm.nih.gov/nuccore/AP011615

13 https://www.ncbi.nlm.nih.gov/nuccore/AC000016.1?fmt_mask=65536

14 https://www.ncbi.nlm.nih.gov/nuccore/BN000507

15 https://www.ncbi.nlm.nih.gov/nuccore/AF055916

16 http://www.uniprot.org/uniprot/P27915

17 https://www.ncbi.nlm.nih.gov/protein/AAG39642

18 https://www.ncbi.nlm.nih.gov/protein/AAG39643

19 ftp://ftp.ncbi.nih.gov/GenBank/gbrel.txt

20 http://www.insdc.org/documents/feature-table

21 https://www.ncbi.nlm.nih.gov/gene

22 https://lucene.apache.org/core/6_1_0/core/org/apache/lucene/search/similarities/TFIDFSimilarity.html

23 http://www.ncbi.nlm.nih.gov/taxonomy

24 http://www.ncbi.nlm.nih.gov/pmc/tools/openftlist/ The version used was downloaded on October 2015.

25 http://lucene.apache.org/

26 The sequences were downloaded on October 2015.

27 https://www.ncbi.nlm.nih.gov/nucest/DW407270

28 The biocuration task carried out in [58] was done for a different context in which analyzing a small sample of records was enough to prove the outcomes claimed in that paper.

29 http://www.ncbi.nlm.nih.gov/pmc/articles/PMC2848993/

30 http://www.ncbi.nlm.nih.gov/nuccore/A23187

31 http://www.ncbi.nlm.nih.gov/nuccore/FJ824848

32 A cloning vector is a small piece of DNA, taken from a virus, a plasmid, or the cell of a higher organism, that can be stably maintained in an organism, and into which a foreign DNA fragment can be inserted for cloning purposes.

33 http://www.ncbi.nlm.nih.gov/pmc/articles/PMC2675058/

34 http://www.ncbi.nlm.nih.gov/nuccore/CP006742

35 http://www.ncbi.nlm.nih.gov/pmc/articles/PMC3923885/

36 http://www.ncbi.nlm.nih.gov/nuccore/BK008760

37 http://www.ncbi.nlm.nih.gov/pmc/articles/PMC4233938/

38 http://www.ncbi.nlm.nih.gov/nuccore/DW407270

39 http://www.ncbi.nlm.nih.gov/pmc/articles/PMC1621083/

## Acknowledgement

The project receives funding from the Australian Research Council through a Discovery Project grant, DP150101550.

## References

[1] Judice L. Y. Koh, Mong Li Lee, and Vladimir Brusic. A classification of biological data artifacts. In Workshop on Database Issues in Biological Databases, pages 53–57, 2005.

[2] Qingyu Chen, Justin Zobel, and Karin Verspoor. Evaluation of a machine learning duplicate detection method for bioinformatics databases. In DTMBIO, pages 4–12, New York, NY, USA, 2015. ACM.

[3] Qingyu Chen, Justin Zobel, and Karin Verspoor. Duplicates, redundancies and inconsistencies in the primary nucleotide databases: a descriptive study. Database, 2017(1):baw163, 2017.

[4] Judice L. Y. Koh, Mong Li Lee, Asif M. Khan, Paul T. J. Tan, and Vladimir Brusic. Duplicate detection in biological data using association rule mining. In European Workshop on Data Mining and Text Mining in Bioinformatics, pages 35–41, 2004.

[5] Steven E. Brenner. Errors in genome annotation. Trends in Genetics, 15(4):132–133, 1999.

[6] Noam Kaplan and Michal Linial. Automatic detection of false annotations via binary property clustering. BMC Bioinformatics, 6(1):1–8, 2005.

[7] Vasilis J. Promponas, Ioannis Iliopoulos, and Christos A. Ouzounis. Annotation inconsistencies beyond sequence similarity-based function prediction – phylogeny and genome structure. Standards in Genomic Sciences, 10(1):108, 2015.

[8] Stephen F. Altschul, Warren Gish, Webb Miller, Eugene W. Myers, and David J. Lipman. Basic local alignment search tool. Journal of Molecular Biology, 215(3):403–410, 1990.

[9] Daniele Apiletti, Giulia Bruno, Elisa Ficarra, and Elena Baralis. Data cleaning and semantic improvement in biological databases. Journal of Integrative Bioinformatics, 3(2):40, 2006.

[10] Alex Rudniy, Min Song, and James Geller. Detecting duplicate biological entities using shortest path edit distance. Int. J. Data Min. Bioinformatics, 4(4):395–410, July 2010.

[11] Min Song and Alex Rudniy. Detecting duplicate biological entities using markov random field-based edit distance. In Bioinformatics and Biomedicine, 2008. BIBM ’08. IEEE International Conference on, pages 457–460, Nov 2008.

[12] Sumithiradevi Chellamuthu and M Punithavalli. Detecting redundancy in biological databases? An efficient approach. Global Journal of Computer Science and Technology, 9(5):141–145, September 2009.

[13] L Holm and C Sander. Removing near-neighbour redundancy from large protein sequence collections. Bioinformatics, 14(5):423–429, 1998.

[14] Weizhong Li and Adam Godzik. Cd-hit: a fast program for clustering and comparing large sets of protein or nucleotide sequences. Bioinformatics, 22(13):1658–1659, 2006.

[15] Eduard Zorita, Pol Cusco, and Guillaume J. Filion. Starcode: sequence clustering based on all-pairs search. Bioinformatics, 31(12):1913–1919, 2015.

[16] Andrew Tritt, Jonathan A Eisen, Marc T Facciotti, and Aaron E Darling. An integrated pipeline for de novo assembly of microbial genomes. PloS one, 7(9):e42304, 2012.

[17] Madison I Dunitz, Jenna M Lang, Guillaume Jospin, Aaron E Darling, Jonathan A Eisen, and David A Coil. Swabs to genomes: a comprehensive workflow. PeerJ, 3:e960, 2015.

[18] Jeroen Crappé, Elvis Ndah, Alexander Koch, Sandra Steyaert, Daria Gawron, Sarah De Keulenaer, Ellen De Meester, Tim De Meyer, Wim Van Criekinge, Petra Van Damme, et al. Proteoformer: deep proteome coverage through ribosome profiling and ms integration. Nucleic acids research, page gku1283, 2014.

[19] Roland J Siezen and Sacha AFT Van Hijum. Genome (re-) annotation and open-source annotation pipelines. Microbial biotechnology, 3(4):362–369, 2010.

[20] Rémi Zallot, Katherine J. Harrison, Bryan Kolaczkowski, and Valérie de Crécy-Lagard. Functional annotations of paralogs: A blessing and a curse. Life, 6(3):39, 2016.

[21] I-Min A Chen, Victor M Markowitz, Ken Chu, Iain Anderson, Konstantinos Mavromatis, Nikos C Kyrpides, and Natalia N Ivanova. Improving microbial genome annotations in an integrated database context. PLoS One, 8(2):e54859, 2013.

[22] Frederic B Bastian, Marcus C Chibucos, Pascale Gaudet, Michelle Giglio, Gemma L Holliday, Hong Huang, Suzanna E Lewis, Anne Niknejad, Sandra Orchard, Sylvain Poux, et al. The confidence information ontology: a step towards a standard for asserting confidence in annotations. Database, 2015:bav043, 2015.

[23] Seán S Óhéigeartaigh, David Armisén, Kevin P Byrne, and Kenneth H Wolfe. SearchDOGS bacteria, software that provides automated identification of potentially missed genes in annotated bacterial genomes. Journal of bacteriology, 196(11):2030–2042, 2014.

[24] Brian P Anton, Simon Kasif, Richard J Roberts, and Martin Steffen. Objective: biochemical function. Frontiers in genetics, 5:210, 2014.

[25] Qingyao Wu, Yunming Ye, Michael K Ng, Shen-Shyang Ho, and Ruichao Shi. Collective prediction of protein functions from protein-protein interaction networks. BMC bioinformatics, 15(2):1, 2014.

[26] Alexandra M. Schnoes, Shoshana D. Brown, Igor Dodevski, and Patricia C. Babbitt. Annotation error in public databases: Misannotation of molecular function in enzyme superfamilies. PLoS Comput Biol, 5(12):1–13, 12 2009.

[27] Friedhelm Pfeiffer and Dieter Oesterhelt. A manual curation strategy to improve genome annotation: Application to a set of haloarchael genomes. Life, 5(2):1427–1444, 2015.

[28] Sylvain Poux, Michele Magrane, Cecilia N Arighi, Alan Bridge, Claire O’Donovan, Kati Laiho, UniProt Consortium, et al. Expert curation in uniprotkb: a case study on dealing with conflicting and erroneous data. Database, 2014:bau016, 2014.

[29] Michael J Bell, Matthew Collison, and Phillip Lord. Can inferred provenance and its visualisation be used to detect erroneous annotation? a case study using uniprotkb. PloS one, 8(10):e75541, 2013.

[30] Maria S Poptsova and J Peter Gogarten. Using comparative genome analysis to identify problems in annotated microbial genomes. Microbiology, 156(7):1909–1917, 2010.

[31] Predrag Radivojac, Wyatt T Clark, Tal Ronnen Oron, Alexandra M Schnoes, Tobias Wittkop, Artem Sokolov, Kiley Graim, Christopher Funk, Karin Verspoor, Asa Ben-Hur, et al. A large-scale evaluation of computational protein function prediction. Nature methods, 10(3):221–227, 2013.

[32] Jesse Gillis and Paul Pavlidis. Characterizing the state of the art in the computational assignment of gene function: lessons from the first critical assessment of functional annotation (cafa). BMC bioinformatics, 14(3):1, 2013.

[33] Indika Kahanda, Christopher S Funk, Fahad Ullah, Karin M Verspoor, and Asa Ben-Hur. A close look at protein function prediction evaluation protocols. GigaScience, 4(1):1, 2015.

[34] EV Koonin and MY Galperin. Sequence-evolution-function: Computational approaches. Comparative Genomics, 2002.

[35] David Lee, Oliver Redfern, and Christine Orengo. Predicting protein function from sequence and structure. Nature Reviews Molecular Cell Biology, 8(12):995–1005, 2007.

[36] Riccardo Percudani, Davide Carnevali, and Vincenzo Puggioni. Ureidoglycolate hydrolase, amidohydrolase, lyase: how errors in biological databases are incorporated in scientific papers and vice versa. Database, 2013:bat071, 2013.

[37] Fenglou Mao, Zhengchang Su, Victor Olman, Phuongan Dam, Zhijie Liu, and Ying Xu. Mapping of orthologous genes in the context of biological pathways: An application of integer programming. Proceedings of the National Academy of Sciences of the United States of America, 103(1):129–134, 2006.

[38] Walter R. Gilks, Benjamin Audit, Daniela De Angelis, Sophia Tsoka, and Christos A. Ouzounis. Modeling the percolation of annotation errors in a database of protein sequences. Bioinformatics, 18(12):1641– 1649, 2002.

[39] Ioannis Iliopoulos, Sophia Tsoka, Miguel A. Andrade, Anton J. Enright, Mark Carroll, Patrick Poullet, Vassilis Promponas, Theodore Liakopoulos, Giorgos Palaios, Claude Pasquier, Stavros Hamodrakas, Javier Tamames, Asutosh T. Yagnik, Anna Tramontano, Damien Devos, Christian Blaschke, Alfonso Valencia, David Brett, David Martin, Christophe Leroy, Isidore Rigoutsos, Chris Sander, and Christos A. Ouzounis. Evaluation of annotation strategies using an entire genome sequence. Bioinformatics, 19(6), 2003.

[40] Judice Lie Yong Koh. Correlation-based methods for biological data cleaning. Master’s thesis, School of Computing National University of Singapore, 2007.

[41] K.N. Srinivasan, P. Gopalakrishnakone, P.T. Tan, K.C. Chew, B. Cheng, R.M. Kini, J.L.Y. Koh, S.H. Seah, and V. Brusic. Scorpion, a molecular database of scorpion toxins. Toxicon, 40(1):23–31, 2002.

[42] Roderic Guigo, Pankaj Agarwal, Josep F. Abril, MoisAõcs Burset, and James W. Fickett. An assessment of gene prediction accuracy in large dna sequences. Genome Research, 10(10):1631–1642, 2000.

[43] G A Seluja, A Farmer, M McLeod, C Harger, and P A Schad. Establishing a method of vector contamination identification in database sequences. Bioinformatics, 15(2):106–110, 1999.

[44] Asif M. Khan, A. T. Heiny, Kenneth X. Lee, Kellathur N. Srinivasan, Tin Wee Tan, J. Thomas August, and Vladimir Brusic. Large-scale analysis of antigenic diversity of t-cell epitopes in dengue virus. BMC Bioinformatics, 7(S-5), 2006.

[45] Kiyoshi Osatomi and Hideo Sumiyoshi. Complete nucleotide sequence of dengue type 3 virus genome RNA. Virology, 176(2):643–647, 1990.

[46] Peter G. Korning, Stefan M. Hebsgaard, Pierre Rouzé, and SÃžren Brunak. Cleaning the genbank arabidopsis thaliana data set. Nucleic Acids Research, 24(2):316–320, 1996.

[47] *The.Gene.Ontology.Consortium*. Gene ontology: tool for the unification of biology. Nat Genet, 25:25–29, 2000.

[48] Database resources of the National Center for Biotechnology Information. Nucleic Acids Research, 44(D1):D7, 2016.

[49] Steve Cronen-Townsend, Yun Zhou, and W. Bruce Croft. Predicting query performance. In Proceedings of the 25th Annual International ACM SIGIR Conference on Research and Development in Information Retrieval, SIGIR ’02, pages 299–306, New York, NY, USA, 2002. ACM.

[50] Ben He and Iadh Ounis. Query performance prediction. Information Systems, 31(7):585–594, 2006. SPIRE 2004(2) Multimedia Databases.

[51] Giridhar Kumaran and Vitor R. Carvalho. Reducing long queries using query quality predictors. SIGIR ’09, pages 564–571, New York, NY, USA, 2009. ACM.

[52] Ben He and Iadh Ounis. Inferring query performance using pre-retrieval predictors. In SPIRE, pages 43–54. Springer Berlin Heidelberg, 2004.

[53] Ying Zhao, Falk Scholer, and Yohannes Tsegay. Effective pre-retrieval query performance prediction using similarity and variability evidence. In 30th European Conference on IR Research, ECIR ’08, pages 52–64, Berlin, Heidelberg, 2008. Springer Berlin Heidelberg.

[54] Stephen E. Robertson, Steve Walker, Susan Jones, Micheline Hancock-Beaulieu, and Mike Gatford. Okapi at trec-2. In TREC, pages 21–34, 1993.

[55] Chengxiang Zhai and John Lafferty. A study of smoothing methods for language models applied to ad hoc information retrieval. In Proceedings of the 24th Annual International ACM SIGIR Conference on Research and Development in Information Retrieval, SIGIR ’01, pages 334–342, New York, NY, USA, 2001. ACM.

[56] Stéphane Clinchant and Eric Gaussier. Information-based models for ad hoc ir. In Proceedings of the 33rd International ACM SIGIR Conference on Research and Development in Information Retrieval, SIGIR ’10, pages 234–241, New York, NY, USA, 2010. ACM.

[57] Eric Sayers. E-utilities quick start. entrez programming utilities help. Technical report, 2010.

[58] Mohamed Reda Bouadjenek, Karin Verspoor, and Justin Zobel. Literature consistency of bioinformatics sequence databases is effective for assessing record quality. Database, 2017(1):bax021, 2017.

[59] Corinna Cortes and Vladimir Vapnik. Support-vector networks. Machine Learning, 20(3):273–297, 1995.

[60] Chih-Chung Chang and Chih-Jen Lin. Libsvm: A library for support vector machines. ACM Trans. Intell. Syst. Technol., 2(3):27:1–27:27, May 2011.

[61] Daniella Schatz, Adva Shemi, Shilo Rosenwasser, Helena Sabanay, Sharon G. Wolf, Shifra Ben-Dor, and Assaf Vardi. Corrigendum. New Phytologist, 206(2):881–881, 2015.

[62] Markus M. Breunig, Hans-Peter Kriegel, Raymond T. Ng, and Jörg Sander. LOF: Identifying density-based local outliers. In Proceedings of the 2000 ACM SIGMOD International Conference on Management of Data, SIGMOD ’00, pages 93–104, New York, NY, USA, 2000. ACM.

